# Inhibitory inputs to avian ITD circuits

**DOI:** 10.1101/2025.11.13.688251

**Authors:** Paula T. Kuokkanen, Z. Mehnoush Faghani, Ira Kraemer, Richard Kempter, Catherine E. Carr

## Abstract

In birds, detection of sound source azimuth begins in the nucleus laminaris (NL), which computes interaural time differences (ITDs). Inhibition has been proposed to protect NL neurons from losing ITD sensitivity with increasing sound level, although the nature of this inhibition is incompletely understood. Anatomical studies show GABAergic synapses throughout NL and the first order nuclei, while *in vitro* studies in chicken NL showed that depolarizing, GABAergic inhibitory postsynaptic potentials (IPSPs) shorten membrane time constants, perhaps to allow the membrane potential to follow rapid synaptic currents accurately over a range of sound levels. Given the importance of inhibition in regulating auditory brainstem activity, we examined the nature of the inhibitory input to NL *in vivo*.

The Superior Olivary Nucleus (SON) is the major source of descending GABAergic inhibition to the ipsilateral nucleus magnocellularis (NM), nucleus angularis (NA) and NL, and receives excitatory input from NA and NL. We used viral tracers to reveal projections from SON to NL, and characterized response types within SON. In NL, we isolated inhibitory synaptic contributions from extracellular field potentials through the iontophoretic application of blockers of GABA (gabazine) or glycine (strychnine). These blockers increase the onset and offset responses evoked by tonal stimuli, but did not shift best ITD. The co-application of gabazine and strychnine revealed supra-linear summation of GABA and glycine. Profiles of synaptic activation revealed more prominent inhibition following stimulus offset, suggesting inputs to NL originate from both the sustained and offset response types of the SON. These heterogeneous responses may represent separate SON neuronal populations.

## Introduction

In birds, detection of sound source azimuth begins with the nucleus laminaris (NL), which computes interaural time differences (ITDs). Inhibition has been proposed to protect NL neurons from losing ITD sensitivity when sound levels increase (Grün et al., 1991; Peña et al., 1996; Brückner and Hyson, 1998; Dasika et al., 2005; Nishino et al., 2008; Burger et al., 2011). This inhibition is assumed to be primarily GABAergic, and to originate from the Superior Olivary Nucleus (SON) (Burger et al., 2011). In both barn owl and chicken, the SON receives excitatory input from nucleus angularis (NA) and NL, and projects to the ipsilateral nucleus magnocellularis (NM), NL and NA (Burger et al., 2005a; Carr et al., 1989; Yang et al., 1999) (Fig. 1). SON has two other major projections; a reciprocal connection to the contralateral SON, and ascending projections to higher-order targets (Conlee and Parks, 1986; Burger et al., 2005a). In chickens, these distinct projections are aligned with three fiber tracts, with each tract originating from specific response types (Baldassano and MacLeod, 2024). Thus, these pathways likely originate from neurons with different *in vivo* response types in the SON, although it remains to be determined which response types project to which nuclei (Coleman et al., 2011; Tabor et al., 2012; Baldassano and MacLeod, 2024; Nishino et al., 2008; Burger et al., 2011).

**Figure 1:**
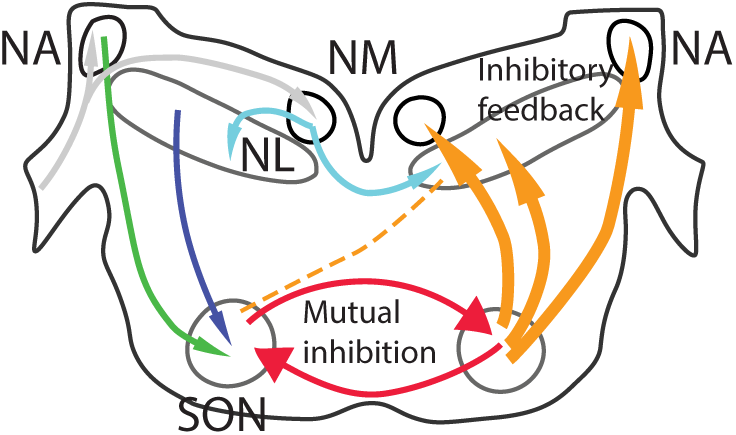
Connections of Auditory Brainstem Nuclei. The Auditory Nerve projects (grey arrows) to Nucleus Angularis (NA) and Nucleus Magnocellularis (NM). NM projects bilaterally (cyan) to Nucleus Laminaris (NL), and both NA and NL project to the SON (blue and green). The SON provides descending inhibition to the auditory brainstem nuclei (orange) as well as mutual inhibition (red).

SON projections subserve distinct functions in NM, NA, and NL. *In vivo*, in chicken NM, activation of SON inputs regulates firing rate, improves phase selectivity; and sharpens frequency tuning (Fukui et al., 2010). These changes are caused by long-lasting, depolarizing IPSPs in NM that are blockable by bicuculline, a GABA_A_ receptor antagonist (Monsivais et al., 2000). In NA, inhibition is both GABAergic and glycinergic (Kuo et al., 2009), and contributes to the intensity coding pathway (MacLeod, 2011). In NL, SON-evoked IPSPs are similarly depolarizing and act to increase low threshold K^+^ conductances and decrease membrane input resistance (Hyson et al., 1995; Funabiki et al., 1998; Brückner and Hyson, 1998). This inhibition reduces the strength of inputs to the NL with increasing stimulus levels (Kuba et al., 2002; Cook et al., 2003; Tang et al., 2009), in part because expression of the low threshold K^+^ conductance reduces the temporal window for summation, effectively increasing the threshold for coincidence of inputs (Hyson et al., 1995; Burger et al., 2011; Tang and Lu, 2012). Thus, inhibition in NL acts to maintain ITD sensitivity by increasing peak-trough contrast. Support for the SON being one source for this inhibition comes from studies of SON lesions in chicken that diminish the contrast between favorable and unfavorable ITDs (Nishino et al., 2008).

Computational models have constrained hypotheses about the role of the SON in ITD tuning. They propose that SON inhibition should be tonic, broadly tuned, and level dependent, and should regulate activity in NM and regulate thresholds in NL (Grün et al., 1991; Peña et al., 1996; Brückner and Hyson, 1998; Dasika et al., 2005; Nishino et al., 2008). In NM, SON inhibition should balance the strength of the inputs from each ear to NM to provide balanced excitation to NL (Burger et al., 2011; Peña et al., 1996; Fukui et al., 2010), achieved by mutual inhibition between SONs (Fig. 1). Thus, a weak, i.e., quiet, input from the ipsilateral ear should provide weak excitation in the ipsilateral SON, which means weak inhibition of inhibitory neurons in the contralateral SON, leading to strong contralateral inhibition (Burger et al., 2011; Fukui et al., 2010). Similarly, strong, i.e., loud, input to the ipsilateral ear should strongly inhibit contralateral inhibitory SON neurons and provide weak inhibitory drive to contralateral NM (Burger et al., 2011; Fukui et al., 2010; Nishino and Ohmori, 2009). In addition to generating balanced inputs to NL, SON inputs within NL should regulate thresholds. ITD selectivity in NL is stable over a range of sound levels in both chickens and barn owls (Peña et al., 1996; Burger et al., 2011; Fukui et al., 2010; Nishino and Ohmori, 2009), with the interrelated mechanisms described above underlying this stability (Kuba et al., 2002; Cook et al., 2003; Agmon-Snir et al., 1998; Reyes et al., 1996; Kuba et al., 2005; Hyson et al., 1995).

Given the importance of inhibition in the regulation of activity in NM and NL (Monsivais and Rubel, 2001; Burger et al., 2011, 2015; Burger and Rubel, 2008; Peña et al., 1996), and especially in understanding the effect of hearing loss on the ITD circuit (Burger et al., 2015; Lu et al., 2018), we have examined the nature of the inhibitory input in NL in the barn owl. The goal of the study was to combine several *in vivo* approaches in the barn owl for comparison with studies in the thoroughly examined chicken model. We used immunohistochemistry, and viral tracing to reveal projections from SON to NL, and characterized response types within the SON. We further combined this work with a study of the iontophoretic application of blockers of GABA or glycine to isolate inhibitory synaptic contributions to extracellular field potentials in NL *in vivo*, similar to the technique used by McColgan et al. (2019).

## Methods

Experiments, including single unit tungsten electrode recordings from SON and multibarrel electrode recordings of the field potential in NL, were conducted at the Department of Biology of the University Maryland. In all, 14 barn owls (*Tyto alba pratincola*) were used to collect the data presented in these and other studies (McColgan et al., 2019; Kraemer and Carr, 2017b). Experimental procedures conformed to NIH Guidelines for Animal Research and were approved by the Animal Care and Use Committee of the University of Maryland. Except for the pharmacology experiments (see below), anaesthesia was induced by intramuscular injections of 10 *−* 20 mg/kg ketamine hydrochloride and 3 *−* 4 mg/kg xylazine. Supplementary doses were administered to maintain a suitable plane of anaesthesia. Body temperature was maintained at 39°C by a feedback-controlled heating blanket.

Recordings were made in a sound-attenuating chamber (IAC, New York). Amplified electrode signals were passed to a threshold discriminator (SD1 (Tucker-Davis Technologies (TDT) Gainesville, FL)) and an analogue-to-digital converter (DD1 (TDT)) connected to a personal computer via an optical interface (OI (TDT)). Acoustic stimuli were digitally generated by custom-made software (‘Xdphys’ written in Dr. Konishi’s lab at Caltech) driving a signal-processing board (DSP2 (TDT)). Acoustic signals were fed to miniature earphones and inserted into the owl’s left and right ear canals, respectively.

### Anatomical studies of SON

At a given recording site in NL, we measured frequency tuning and tuning to ITD followed by microinjection of 50 *−* 100 nL virus-containing rAAV2-retro tdTomato solution (Tervo et al., 2016). After a survival time of 2-3 weeks, owls were perfused with 4% paraformaldehyde, and brains blocked and sectioned. Sections were examined using confocal microscopy and reconstructed with the aid of a Neurolucida system.

For immunohistochemical studies of GABAergic terminals, owls were perfused as above, and sectioned at 30 µm. The antiserum against Vesicular Glutamate Transporter (VGAT) was obtained from Antibodies Inc (Davis, CA). Free-floating sections were incubated for 48 h at 4°C in primary antiserum diluted to 1:1000 in PBS containing 0.5% Triton X, followed by 24 h at 4°C in biotinylated secondary antiserum. Sections were subsequently treated according to the avidin-biotin method (Vector laboratories, Newark, CA). Additional immunohistochemical material included material incubated with antisera against Glutamic Acid Decarboxylase (GAD) (Carr et al., 1989) and material kindly provided by Winer and LaRue, which used rabbit-anti-glycine antiserum (courtesy of R.J. Wenthold; Gly II, 1:1,500), or rabbit-anti-GABA antiserum (from R.J. Wenthold; 1:2,000) (Winer et al., 1995). Material was analyzed using the Neurolucida system (Microbrightfield, VT, USA).

The immunohistochemically labeled SON neurons were categorized by shape into two classes: fusiform cells and multipolar cells, such that a cell was classified as fusiform if both the (Neurolucida parameter) ‘roundness’ of the cell was *<* 0.65 and the ‘formfactor’ was *<* 0.83.

### Pharmacology

We used iontophoresis to deliver drugs to NL (McColgan et al., 2019). All data were obtained using glass multibarrel electrodes (Carbostar-3 or -4 LT, Kation Scientific, Minneapolis, MN). Microelectrode barrels were filled with gabazine (3 mM SR95531), a GABA_A_ receptor antagonist and/or strychnine (10 mM), a glycine receptor antagonist (Sigma, St. Louis, MO). An additional control experiment used saline alone. After control data were collected in the beginning of each experiment, drugs were ejected for 15 *−* 20 min, followed by a 20 min washout period, then the cycle repeated. Analyses revealed that repeated washouts did not always return to baseline, especially with strychnine. Thus, we used only the initial condition as control. These are terminal experiments that use 8 –10 ml/kg of 20% urethane urethane as an anesthetic (McColgan et al., 2019).

### In vivo physiology in SON

For these experiments, we recorded in SON with tungsten electrodes (5 *−* 20 MΩ). A grounded silver chloride pellet served as a reference. We measured monaural and binaural frequency tuning, tuning to ITD, and responses to changes in level. In some cases, lesions were used to confirm the location of recordings. We used WaveClus MATLAB package for the spike sorting of the waveforms (Quian Quiroga et al., 2004).

### Data collection and analysis of extracellular field potentials in NL

In each recording location, we measured monaural and binaural frequency tuning in the control condition (best frequencies (BF) of the neurophonic ranging 3.2 to 6.8 kHz). For each drug condition (including control conditions), we measured responses to binaural 100-500 ms duration tone-pips at BF typically at 40 dB SPL, as well as 1-2 additional frequencies around BF, the respective ITD tuning curves (minimum: favourable and unfavourable ITDs) and one or more stimulus amplitudes.

Data recorded during iontophoresis were analyzed using a custom library (pyXdPhys, https://github.com/phreeza/pyXdPhys) by McColgan et al. (2019). Response traces from each drug condition were averaged and filtered (low-pass: *<*1 kHz, high-pass: *>*1.5 kHz). The amplitudes of the low pass filtered onsets (offsets) were calculated from the 0 *−* 50 ms time span after the stimulus onset (offset) and normalized by the respective control condition onset (offset).

We note that over the course of hours, overlapping but not coinciding with the administration of drugs and/or saline, there were small (typically some tens of degrees, corresponding to roughly half of it in microseconds) inconsistent phase shifts in the field potential at the stimulus frequency (neurophonic). For such shifts *>* 20 degrees, the phase shifts of the monaural responses were of opposing directions. The phase shifts were not systematically correlated with either the low-pass or the high-pass response amplitudes across the owls (Pearson correlation coefficients for LP: mean *±* SD = 0.0 *±* 0.4, ranging from *−*0.47 to 0.52; for HP 0.0 *±* 0.4, ranging from *−*0.71 to 0.62, with similar variability in all drug groups). These occasional phase shifts may be associated with spatial drift of the electrode with respect to the brain tissue over hours, consistent with the opposite changes in monaural neurophonic phase shifts.

## Results

To examine the nature of the inhibitory input from SON to NL, we used immunohistochemical techniques to label neurons in SON and viral tracers to reveal projections between NL and SON. We also characterized *in vivo* neuronal responses in the SON. We then used iontophoretic application of GABA and glycine blockers *in vivo* into NL, to isolate inhibitory synaptic contributions in NL.

### SON contains neurons immunopositive for GABA and glycine

The SON is located in the rostral pons, ventral to NL (Fig. 1). It contains multiple cell types, some of which are densely labeled by antisera directed against GABA or GAD (Carr et al., 1989; Code et al., 1989; Lachica et al., 1994; Von Bartheld et al., 1989). We measured labeled cells, and found based on the shape of the neurons two sizes of GAD+ neurons: large multipolar cells (cell body area of about (mean *±* SD): 371 *±* 147 *µ*m^2^, *N* = 52) and medium-sized fusiform cells (cell body area of about 229 *±* 69 *µ*m^2^, *N* = 49) (Fig. 2A); and there is a third group of large unstained neurons (Carr et al., 1989). The SON also contains a similar population of neurons labeled by antisera against glycine in chicken (Coleman et al., 2011) and in barn owl (Fig. 2B). Again, there were two sizes of glycine+ neurons: large multipolar cells (cell body area of about 384 *±* 125 *µ*m^2^, *N* = 53) and medium-sized fusiform cells (cell body area of about 199 *±* 85 *µ*m^2^, *N* = 56) (Fig. 2B). The sizes of the neurons in each size group were not significantly different between the GAD+ and glycine+ populations when corrected for multiple comparisons (Student’s 2-population tests, Šidák-correction for *p* = 0.05, Abdi, 2007).

**Figure 2:**
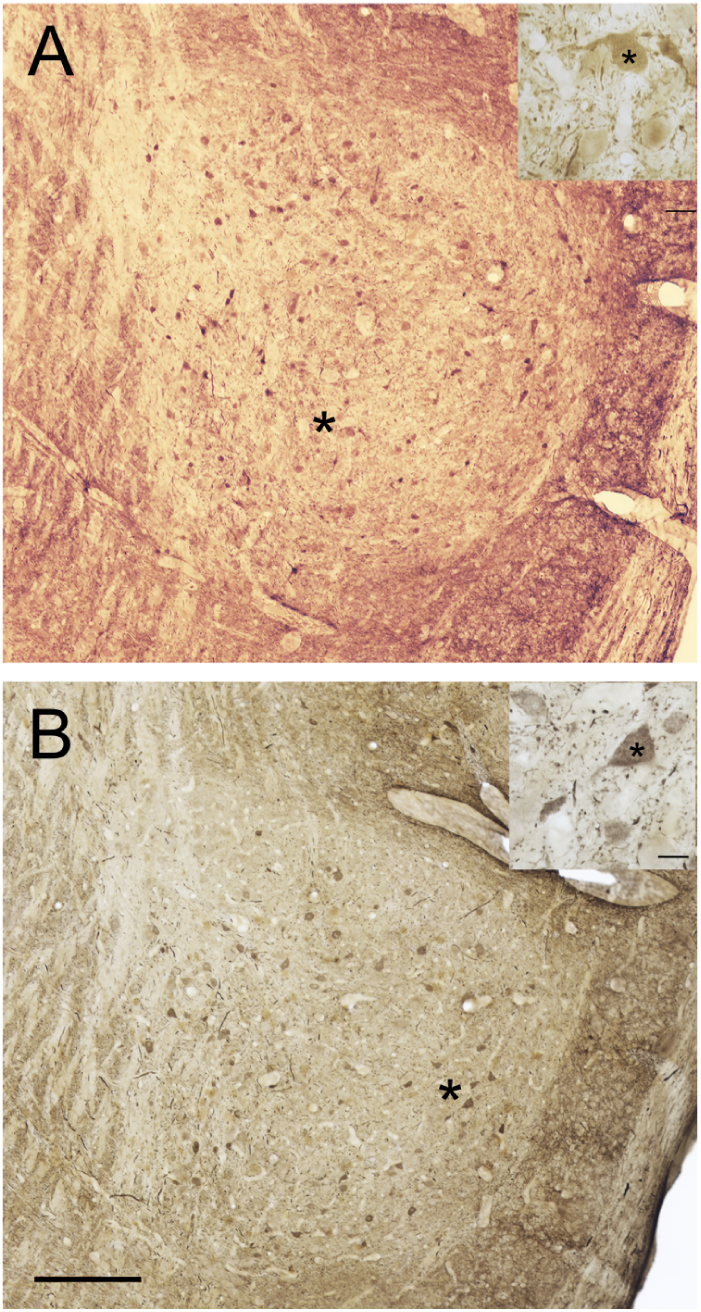
Immunoreactivity in SON. A. Anti-GABA immunoreactive neurons in a transverse section through SON. Inset: immunoreactive neuron (*) surrounded by immunoreactive terminals and unlabeled cell bodies. B: Anti-Glycine immunoreactive neurons in SON. Inset: immunoreactive neuron (*) surrounded by immunoreactive terminals and unlabeled cell bodies. Location of inserts marked by * next to the cell body. Scale bar for A and B = 200 *µ*m, and for both insets = 20 *µ*m.

Because we did not double label SON cells, we were unable to determine whether SON neurons co-expressed GABA and glycine. Nevertheless, our data suggests co-expression because the labeled neurons fell into two similar size classes for both labeling methods (see above). Furthermore, immunohistochemical evidence has suggested that GABA and glycine may be co-released at SON target neurons, and glycinergic responses have been shown in NA, NL, and SON (Kuo et al., 2009; Fischl et al., 2014; Coleman et al., 2011; Fischl and Burger, 2014).

### Connections of SON

Many GABAergic and glycinergic immunolabeled neurons in SON are likely to project to NA, NL, and NM, and form perisomatic terminals in NL (see the next section) and NM (Code et al., 1989; Lachica et al., 1994; Coleman et al., 2011; Nishino et al., 2008; Fukui et al., 2010). However, we note that the two SON are also reciprocally connected, and Burger et al. (2011) have proposed that this contralateral projecting pathway is inhibitory. Thus, we cannot assume that all inhibitory neurons project to NM and back to NA and NL.

SON has been shown to project ipsilaterally to NL in the barn owl (Takahashi and Konishi, 1988). To identify which SON neurons projected to NL, we injected rAAV2 into NL. Injections in three barn owls confirmed that SON projects ipsilaterally to NL. A sample of retrogradely labeled neurons in SON revealed medium-sized and large-sized neurons (Fig. 3A). The sizes of neurons in the medium-sized and the large-sized populations were not significantly different from the GAD+ populations or glycine+ populations described in the previous section (all *p >* 0.2, Student’s 2-population tests). Although rAAV2 principally retrogradely labeled SON neurons, note that Fig. 3B also shows anterogradely labeled NL axons both passing through and terminating in SON. In addition to retrogradely labeled neurons in the ipsilateral SON, we also found a small number of labeled neurons in the contralateral SON (not shown). We therefore indicate in Figure 1 (dashed line) that the barn owl NL receives a weaker projection from the contralateral SON.

**Figure 3:**
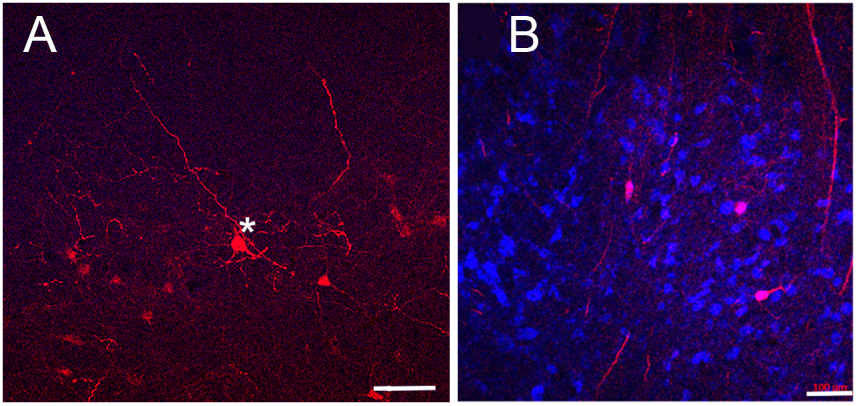
SON Projection Neurons. A. Injection of rAAV2-retro tdTomato into NL yielded two morphological types of projection neuron in SON, of large (*) and medium size. B. rAAV2-retro labeled SON neurons with labeled NL axons. Blue neurotrace counterstain. Scale bars=100 *µ*m

### GABAergic and glycinergic terminals in NL

The projection from the SON to NL is marked by an oblique lateral to medial axon tract that descends from SON to NL, composed of fine axons that project into NL. Each NL neuron is surrounded by a network of terminals, which can be labeled by antibodies against VGAT. VGAT labels both glycinergic and GABAergic terminals (MacLeod and Pandya, 2022) (Fig. 4).

**Figure 4:**
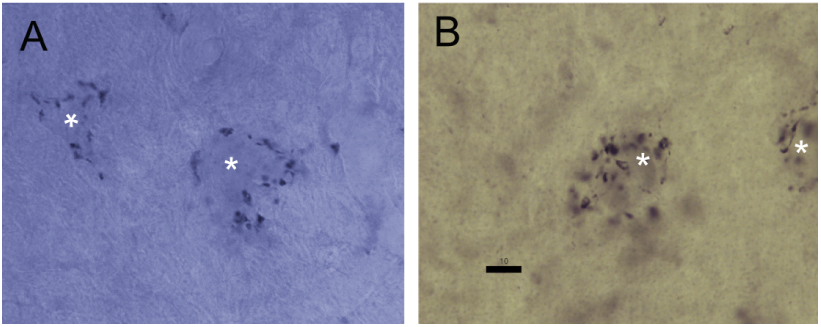
GABAergic Terminals in NL. A. NL neurons (*) surrounded by perisomatic VGAT+ terminals. B. NL neurons (*) surrounded by perisomatic GAD+ terminals. Scale bar for both=10 µm.

GABAergic terminals may also be labeled with antisera against GAD. Comparisons of GAD and VGAT material revealed similar numbers of terminals per cell. Since NL neurons have such short dendrites that their dendritic fields do not overlap (Carr and Boudreau, 1993), we were able to count the number of terminals per cell. Each NL neuron was surrounded by about 53 *±* 14 (mean *±* SD, *n* = 80) GAD+ and 58 *±* 17 (*n* = 96) VGAT+ terminals. By contrast, the small low best frequency region (*<* 1 kHz) has a higher terminal density (Carr et al., 1989; Nishino et al., 2008).

### In vivo physiology in SON

The SON is morphologically heterogeneous and projects to multiple targets (Figs. 1–3) (Takahashi and Konishi, 1988; Baldassano and MacLeod, 2024). Given the heterogeneity of the SON cell types and the array of response types found in chicken (Tabor et al., 2012; Lachica et al., 1994), barn owl SON should also contain multiple response types. To test this hypothesis, we characterized barn owl SON responses *in vivo* in seven owls. Lesions were used in some cases to confirm recording locations (Fig. 5D). We isolated 70 single units that responded to auditory stimuli, and report upon a portion of the data here. Most units (65 out of 70) responded preferentially to binaural noise and showed broad frequency tuning with low firing rates (Kraemer and Carr, 2017a,b).

**Figure 5:**
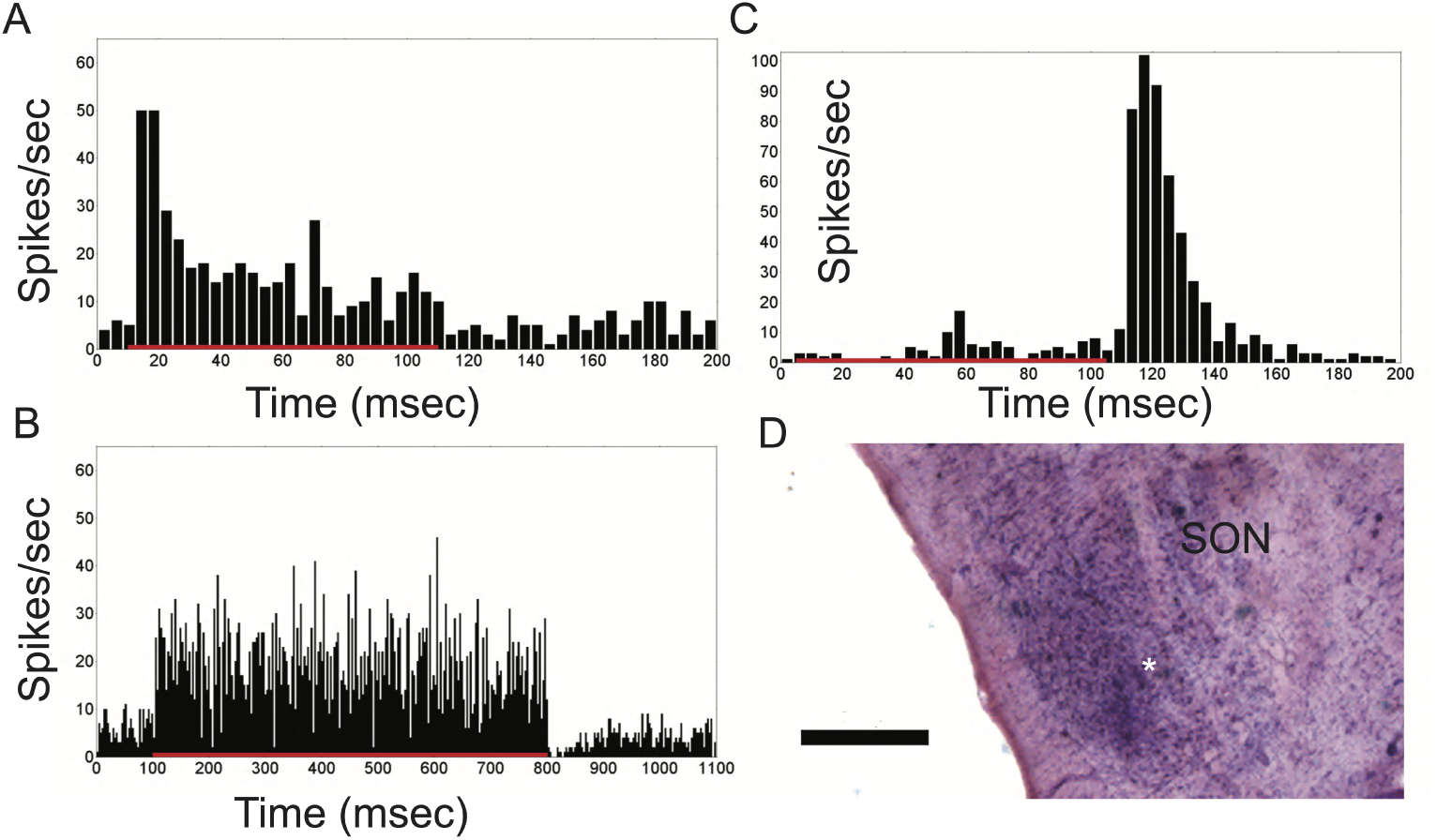
SON Response Types. A. Peristimulus time histogram (PSTH) of a cell with a primary-like response, with an initial peak followed by tonic firing throughout the duration of the stimulus (10 *−* 110 ms, red bar). B. PSTH of a cell with a sustained response, which typically fires throughout the stimulus. C. PSTH of an offset response cell, which has a high response following the stimulus. D. Following several experiments, recording sites were confirmed by the presence of a lesion from the recording electrode (*). Scale bar=500 µm.

The SON included several response types: sustained, primary-like, off, onset, and on-off. Sustained responses, including primary-like responses (Fig. 5A,B), were the most common (70%, *n* = 50), followed by 13% off responses (Fig. 5C) and altogether 17% onset and on-off responses (not shown). There was potential for separating sustained response types into primary-like and sustained (Fig. 5A, B), but these two response types were grouped because peristimulus time histogram (PSTH) shapes revealed a continuous distribution. Off-response units were easy to separate because they fired vigorously after the stimulus ended (Fig. 5C). Based on the shape of the rate-level function of binaural noise stimuli from 15 *−* 70 dB SPL, most units demonstrated monotonic rate level curves, and most were excited by both ears (Kraemer and Carr, 2017a,b). Only about 21% of the 70 SON recordings were ITD sensitive, with ITD sensitivity distributed among all response types.

### Pharmacology in NL

*In vitro* studies in chicks had shown presynaptic GABA release and postsynaptic receptor subtypes in NL (Tang and Lu, 2012), with activity-dependent responses to sensory manipulations (Lu et al., 2018; Fischl and Burger, 2014; Fischl et al., 2014). To examine the role of GABAergic innervation in the NL of owls, we isolated inhibitory synaptic contributions from the extracellular field potential through the iontophoretic application of blockers of GABA_A_ receptors. This approach is similar to our previous analysis of AMPA sensitive responses in NL (McColgan et al., 2019). In that study, we isolated *in vivo* AMPA sensitive synaptic contributions from the extracellular field potential (EFP) in NL (McColgan et al., 2019) using iontophoretic application of the antagonist NBQX (Kuba et al., 2002; Cook et al., 2003).

Here, we analyzed EFPs in NL during tone-burst stimuli averaged over trials in the control condition (Fig. 6A, blue) and during gabazine application (Fig. 6B orange, overlap brown). In low-pass filtered responses (Fig. 6Aii,Bii), we subtracted gabazine responses from control conditions to reveal synaptic contributions to the EFP (Fig. 6Biii). We found that upon iontophoresis of gabazine, low-pass filtered responses increased in amplitude, consistent with blockade of inhibitory conductances with respect to the control condition. This relative increase in the EFP amplitude with the postsynaptic blockage of chloride channels is in line with reduction of an outward current. The example in Fig. 6E shows that the standard deviation of the low-pass filtered responses during the sustained activity was strongly affected by the drug condition. In contrast, there was no significant change in the amplitude of the high-pass filtered response (Fig. 6Ai,Bi,E): the standard deviation of the high-pass filtered sustained component (*>* 1 kHz, see also ‘*Drug-insensitivity of extracellular ITD tuning*’) remained unchanged relative to the initial control condition. Our results also align with *in vitro* studies in chick (Burger et al., 2011) and *in vivo* and theoretical studies in owl (Peña et al., 1996).

**Figure 6:**
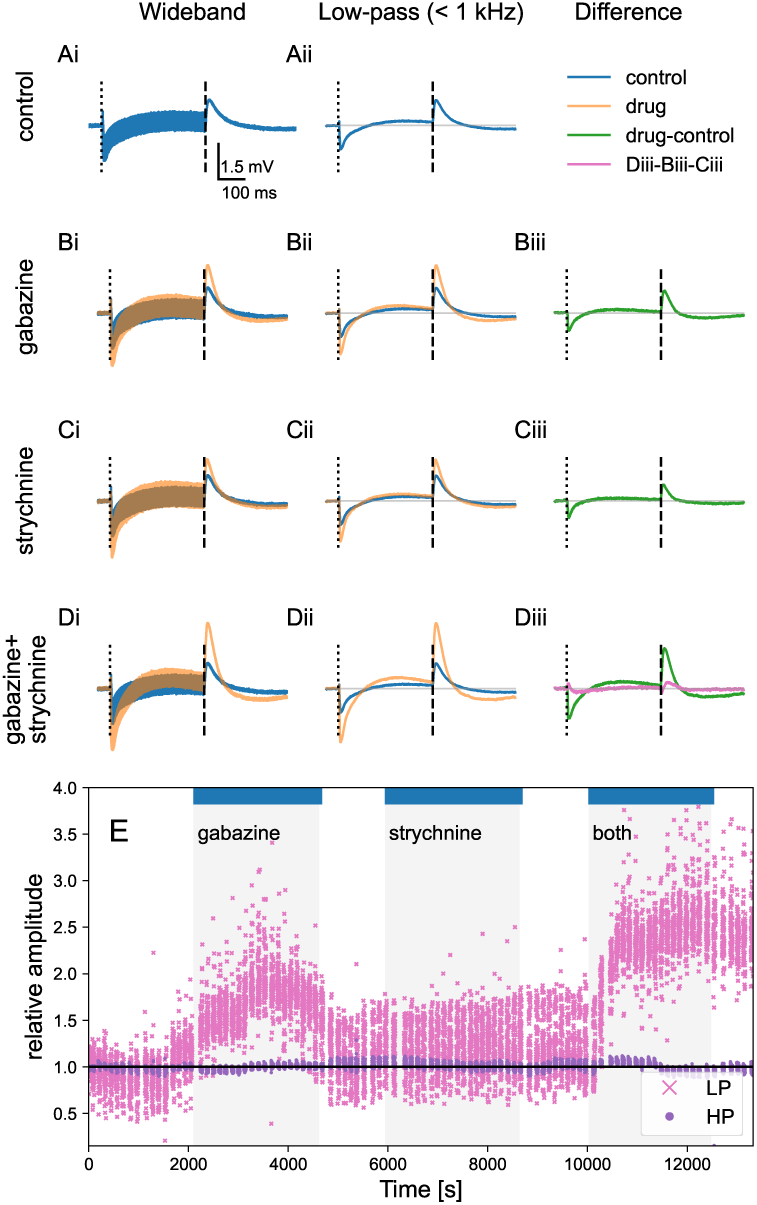
Separability of Synaptic Contributions. Gabazine iontophoresis alters extracellular field potentials in NL, with averaged low-pass filtered responses in control (blue) and during drug application (orange; overlap in brown). Stimulus frequency 5600 Hz, amplitude 40 dB, ITD: 30 *µ*s. Stimulus onset (50 ms): dashed lines in A-D, stimulus offset (450 ms): dash-dotted lines in A-D. **A.** Control condition: i) wide-band response, ii) low-pass filtered response (*<* 1 kHz). **B.** Gabazine, i-ii) control (blue) and drug condition (orange) shown; overlap in brown; iii) difference to control condition for the low-pass filtered responses (green). **C.** Strychnine. **D.** Gabazine and strychnine applied simultaneously. Diii) Pink: the difference of the synaptic contributions when both drugs applied simultaneously (green line in Diii) and the sum of separately applied drugs (Biii and Ciii) reveals the supra-linear effect of GABA and glycine. **E.** The response amplitude of the high-pass (HP, purple) filtered sustained response (standard deviation of the neurophonic 50-400 ms after stimulus onset) and of the low-pass (LP, pink) filtered onsets (average of the trace 0-50 ms after stimulus onset). All stimulus conditions during the experiment are shown (amplitudes: 30 and 40 dB SPL, stimulus frequencies: 5.3 and 5.6 kHz, ITDs: *−*30 and 60*µ*s, resulting in 8 different stimulus combinations), each normalized with the average respective response in the initial control condition with no iontophoresis (0-2100 sec).

*In vitro* data support co-release of GABA and glycine from SON terminals in target nuclei, with physiological responses to glycine shown in NA, NM, NL and SON (Kuo et al., 2009; Coleman et al., 2011; Fischl et al., 2014). We isolated glycine-sensitive synaptic contributions from the field potential in NL *in vivo* using iontophoretic application of the antagonist strychnine. Similar to the gabazine condition, we compared strychnine responses with the control condition (Fig. 6C). Strychnine iontophoresis was associated with an increase in the amplitude of the low-pass filtered responses at both onset and offset amplitude, for all of the stimulus conditions. In the example shown in Fig. 6C,E, the increase with the strychnine iontophoresis, i.e., the glycinergic postsynaptic contribution, was atypically small (see also Fig. 7A,B,Cii).

**Figure 7:**
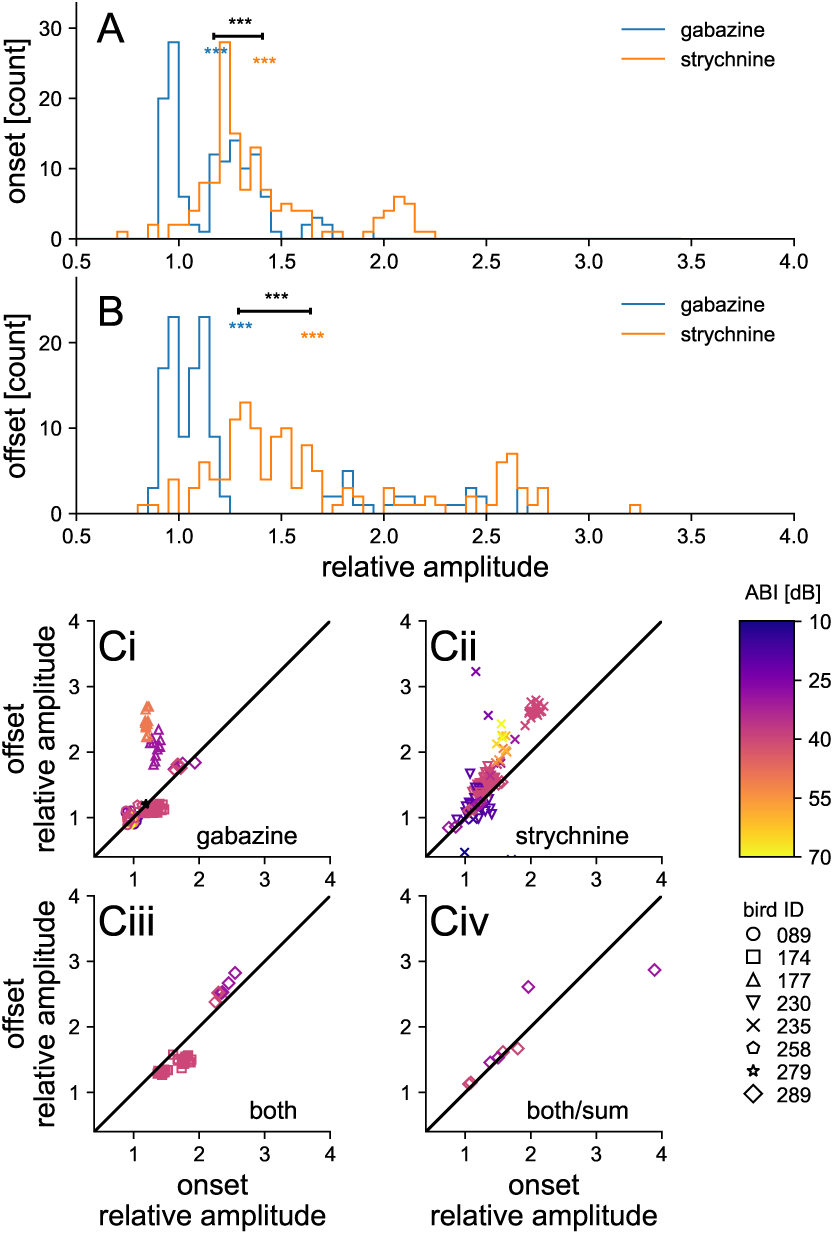
Population Responses to Pharmacology. Across the population, both gabazine and strychnine application significantly increased the onset and the offset responses with respect to the control condition. The response was measured as the average of first 50 ms of the low-pass filtered EFP after the stimulus onset/offset, and averaged across all ITDs. **A.** Onset response, ratio to control condition. Gabazine: mean *±* SE: 1.17 *±* 0.02, *p* = 3 *·* 10*^−^*^15^, *N* = 133. Strychnine: 1.41 *±* 0.03, *p* = 6 *·* 10*^−^*^30^, *N* = 137. The effect of strychnine was larger than that of gabazine (*p* = 2 *·* 10*^−^*^11^, Student’s 2 population test). **B.** Offset response, ratio to control condition. Gabazine: 1.29 *±* 0.05, *p* = 3 *·* 10*^−^*^10^, *N* = 133. Strychnine: 1.64 *±* 0.05, *p* = 4 *·* 10*^−^*^27^, *N* = 137. Also for the offset, the effect of strychnine was larger than that of gabazine (*p* = 9 *·* 10*^−^*^8^). **Ci-Ciii.** Relative increase of the onset response (x-axis) and the offset response (y-axis) for different drug conditions and for the sum of gabazine and strychnine when measured independently in the same recording location. For both gabazine and strychnine, the increase of the offset was larger than that of the onset (*p* = 0.0010 and *p* = 3 *·* 10*^−^*^5^, respectively). Stimulus amplitude is shown with the color scale. The stimulus frequency, spanning from 3.2 kHz to 6.8 kHz, did not systematically affect the relative increase. **Civ.** Non-linearity of GABA and glycine effects: for simultaneous administration of gabazine and strychnine (‘both’) the effect on both onset and offset was larger than expected from the ‘sum’ of both drugs administered independently (both/sum *>* 1). Note that the experiment including both separate and simultaneous administration of drugs was only possible in a single animal due to its duration.

We also studied the interaction of the two drugs (Fig. 6D). Simultaneous injection of gabazine and strychnine revealed a synaptic interaction component when compared to the sum of independent injections (Fig. 6Diii, pink line). The interaction component was larger than the sum, with a longer duration at the tone offset compared to the onset; see also Fig. 6E, right.

#### Differential effects of gabazine and strychnine

For iontophoresis of either drug, the effect of the strychnine in the population were typically larger than that of gabazine. Both drugs had a larger effect at tone offset than at tone onset (Fig. 7A,B), and simultaneous drug application had a larger effect than expected from the sum of independent drug applications (Fig. 7Civ).

The absolute amplitude of the onset and offset responses, i.e., of the synaptic contributions, increased with level, while their relative amplitude in comparison with the respective control condition remained roughly level-independent (Fig. 7Ci,Cii) in the birds in which we recorded a broad amplitude range. Between experiments, the time courses of the onset and offset varied in both their duration and relative amplitude. Modeling studies suggest that differences in electrode position with respect to the cell body may underlie variation in extracellular synaptic potential recordings (McColgan et al., 2019).

#### Drug-insensitivity of extracellular ITD tuning

Inhibition in ITD coding has been the subject of numerous studies (for reviews, see Roberts et al., 2013; Yin et al., 2019). However, it has been difficult to examine the effects of GABA on ITD sensitivity *in vivo* in NL (Takahashi and Konishi, 2002). Thus, here we again compared the control and drug responses in NL for tone-burst stimuli. We again study the impact on the low-frequency responses that we had already characterized (Fig. 8A,B) but now also focus on the high-frequency parts of the responses (Fig. 8C,D).

**Figure 8:**
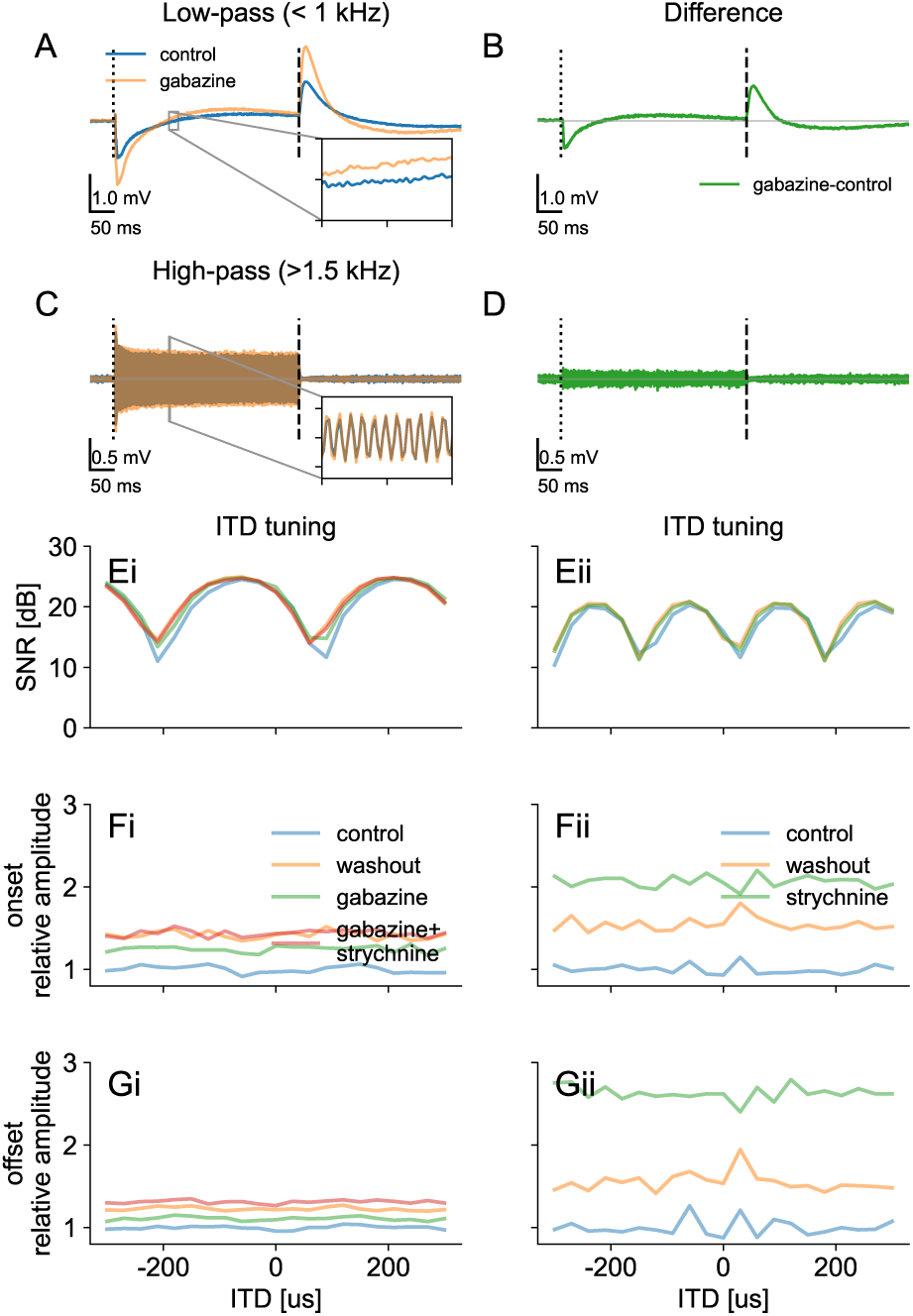
ITD Tuning of the Neurophonic was not Altered by Gabazine or Strychnine. **A-D** Separation of the EFP response to low- and high-pass filtered components (same owl as in Fig. 6). **A.** Low-pass (*<* 1 kHz) filtered EFP responses. Inset: a 10-ms (0.5-mV) excerpt for detail. **B.** Difference between the low-pass filtered responses in drug and control conditions reveals synaptic contributions. **C.** High-pass (*>* 1.5 kHz) filtered EFP responses. There is no change in the envelope amplitude, but a phase shift of 18 degrees of the neurophonic (corresponding to *−*9 *µ*s for the stimulus frequency of 5600 Hz). Inset: a 1-ms (1.5-mV) **D.** Difference between the high-pass filtered responses in drug and control conditions. Because the nearly-sinusoidal traces have a slight respective phase shift, their difference has a non-zero amplitude. The constant envelope amplitude of the difference is consistent with a constant phase shift between the traces. **E–G** Representative examples of recordings with gabazine and strychnine from two different birds (Ei and Eii). **E.** The signal-to-noise ratio of the neurophonic, i.e. spectral peak at the stimulus frequency wrt. background spectrum (Kuokkanen et al., 2010), showed strong ITD tuning that was not systematically modulated by the application of synaptic blockers. **F,G.** The low-pass filtered onset (F) and offset (G) were increased both by gabazine (i) and by strychnine (ii) application but not ITD-modulated. The values in F,G are normalized by the average of the respective control condition across all ITDs.

We found that the ITD tuning of the high-frequency EFP (‘neurophonic’) (Fig. 8C,E) was not affected by either drug. The neurophonic is composed almost exclusively of the incoming axonal spikes (see Discussion; Kuokkanen et al., 2010, 2013), and was not expected to change with modified inhibitory synaptic contributions. In most experiments, there were small (typically 10–30 degrees at the best ITD) drifts in the neurophonic oscillation phase (Fig. 8C, inset) during the hours of recording time. These shifts were possibly related to changes of the location of the electrode with respect to the tissue, i.e., the delay line axons. These drifts typically did not coincide with the time of the drug injections, but were consistent with the small shifts in the neurophonic ITD tuning (e.g. Fig. 8E) (Kuokkanen et al., 2013).

The inhibitory synaptic components of the field potential, i.e. the onset and offset of the low-pass filtered response, as revealed by the drugs, were not ITD tuned (Fig. 8F,G). Observing no ITD tuning also supported our hypothesis that this low-pass EFP did not include contributions from the postsynaptic NL neurons that are strongly ITD tuned (Peña et al., 1996; Funabiki et al., 2011, see also Discussion). We note that for the population data shown in Fig. 7, we took advantage of the ITD insensitivity of the onset and offset responses, which justified there the averaging the responses across all ITDs for more robust results.

The finding that inhibitory synaptic components in NL are not ITD tuned is consistent with our SON recordings, where only a low fraction of about 21% of the SON neurons were ITD tuned (Kraemer and Carr, 2017a). Furthermore, the SON projections into NL appear widespread (Nishino et al., 2008), which is also consistent with the imprecise mapping of ITD.

## Discussion

GABAergic terminals surround all NL neurons (Carr et al., 1989). These inhibitory inputs should protect NL neurons from losing their ITD sensitivity when sound levels increase (Grün et al., 1991; Peña et al., 1996). To examine the role of inhibition in encoding ITD, we isolated inhibitory synaptic contributions from extracellular field potentials in the NL of barn owls through the iontophoretic application of blockers of GABA or glycine. Both blockers caused significant increases in the low spectral frequencies of the field potential that are associated with synaptic currents (Kuokkanen et al., 2018; McColgan et al., 2019), but did not affect the ITD tuning of the neurophonic responses in NL. Although we were unable to identify which SON response types projected to NL, synaptic activation profiles, in conjunction with previous modeling work of the excitatory synaptic currents (McColgan et al., 2019), suggest that the inhibitory inputs to NL could originate from several SON response types.

### The role of inhibition in ITD coding

Inhibition in ITD coding has been the subject of extensive research (for reviews see Roberts et al., 2013; Yin et al., 2019). In small mammals, manipulation of glycinergic inhibition affects the best ITD of responses in the medial superior olive (Franken et al., 2015; Leibold and Grothe, 2015) in a physiologically appropriate fashion. In birds, however, GABAergic inhibition in NL appears instead to increase the dynamic range of ITD sensitive responses in NL (Grün et al., 1991; Peña et al., 1996; Nishino et al., 2008) and not alter ITD tuning. *In vitro* studies further support a direct effect of inhibition on coincidence detection (Brückner and Hyson, 1998; Funabiki et al., 1998), where inhibition speeds the kinetics of excitatory postsynaptic potentials, thus reducing the duration of the coincidence window.

For our study, it is likely that the iontophoretic application of gabazine in NL blocked GABA_A_ inhibition directly and locally, leading to the observed increases in field potential (Fig. 7). The local block probably dominates the indirect effects from the SON, although the different roles for inhibitory activity discussed below must be considered in a system with activity-dependent feedback. If the barn owl circuit resembles the chicken, the negatively coupled SONs should interact and generate inhibitory feedback bilaterally that offsets imbalances in acoustic intensity at the two ears, such as those generated by unilateral hearing loss (Burger et al., 2011; Fukui et al., 2010).

In NL, GABA_A_ signaling is depolarizing (Hyson et al., 1995). The GABAergic input activates low threshold K+ conductances to decrease the membrane time constant and improve coincidence detection (Ashida et al., 2007; Kuba et al., 2003, 2005). Chicken NL also contains abundant GABA_B_ receptors (Burger et al., 2005b; Tang et al., 2009), shown to have a largely presynaptic action (Tang et al., 2009). Future studies might address the complex interactions among GABA_B_ and metabotropic glutamate receptors *in vivo*.

The avian auditory brainstem nuclei exhibit a diversity of inhibitory transmission that suggests differential glycine, GABA_A_ and GABA_B_ receptor activity, tailored to roles in NM, NL, and NA (Goel et al., 2020). These distinct roles are supported by the presence of divergent inputs from the SON (Kuo et al., 2009; Baldassano and MacLeod, 2024; Burger et al., 2005a). Furthermore, it seems likely that glycine may be co-released with GABA (Kuo et al., 2009; Coleman et al., 2011; Fischl et al., 2014). These *in vitro* results were supported by our iontophoresis of strychnine, which on its own typically did not affect the recorded potential in NL, but revealed a pronounced synergy when iontophoresed with gabazine (Fig. 7).

Results from our study suggest that neither GABAergic nor glycinergic inhibition shifts the ITD tuning of the NL neurons’ activity: there were no systematic phase shifts of the neurophonic phase associated with the drugs, and the inhibitory synaptic contributions were not ITD tuned (Fig. 8F,G). Instead, the effects of synaptic blockers support the observations of Peña et al. (1996) and are consistent with providing an increased dynamic range. We also observed varying profiles of synaptic activation, consistent with inputs from different SON cell types. Profiles of synaptic activation revealed more prominent inhibition following stimulus offset than onset (Fig. 7), suggesting inputs to NL might originate from both the primary-like, on-off and offset response types of the SON. These heterogeneous synaptic responses may represent separate SON neuronal populations and allow for additional regulation of ITD coding. It remains to be tested whether the ITD tuning of postsynaptic NL neurons could be modified by inhibition: showing postsynaptic ITD modulation in the EFP remains challenging (Kuokkanen et al., 2013, 2018) and was not possible in our data set (see below). It is likely that postsynaptic ITD tuning isunaffected by inhibition, since a prior study that combined iontophoresis of GABA or muscimol (a GABA_A_ receptor agonist) in NL with recordings in the optic tectum did not show changes in best ITD (Takahashi and Konishi, 2002).

### In vivo physiology in SON

The projections of the SON in the barn owl resemble those in chicken (Nishino et al., 2008; Von Bartheld et al., 1989; Coleman et al., 2011) and zebra finch (Wild et al., 2010). Similarly, most of the units we recorded in SON were similar to those recorded in chicken, despite the use of either monaural or free field stimulation in chicken (Tabor et al., 2012; Lachica et al., 1994). In our sample, SON responses were largely binaural, and characterized by a majority of sustained response units, with a smaller number suppressed by sound. A previous study in barn owl (Moiseff and Konishi, 1983) had found both monaural and binaural responses, consistent with direct inputs from the ipsilateral NA and NL (Takahashi and Konishi, 1988). The heterogeneous response types may represent separate SON neuronal populations projecting to different nuclei (Burger et al., 2005a; Baldassano and MacLeod, 2024).

### Components of the extracellular field in NL

Isolating single spikes from the extracellular field potential (EFP) in NL is challenging because they are masked by the millivolt-scale neurophonic potential (Funabiki et al., 2011). The neurophonic potential follows the temporal features of the stimulus and reflects the summation of the recorded phase-locked population activity. Most of the high-frequency component of the EFP in NL originates from the incoming axonal spikes of NM axons (Kuokkanen et al., 2010, 2013, 2018). Hundreds of axons in the vicinity of the recording electrode conduct the inputs to the NL neurons, phase-locked up to BFs of 9 kHz (Köppl, 1997a,b). Both for NL neurons, and onto the recording electrode, phase-locked inputs summate: For an NL neuron, coincident synaptic inputs summate (Ashida et al., 2012) to elicit postsynaptic spikes that in turn map the ITD of the ipsi- and contralateral inputs. By contrast to the synaptic contributions that are strongest around the cell bodies, axonal contributions from the nodes of Ranvier are more evenly distributed across the tissue and thus picked up by the electrode (Kuokkanen et al., 2010, 2013). The axonal spikes have a temporally narrow waveform, contributing to the EFP at mostly at 2-8 kHz spectral frequencies. As expected, these axonal spikes have a shorter duration than the post-synaptic spikes, which mostly contribute to the 300-1500 Hz frequency band of the spectrum (Kuokkanen et al., 2018). This spectral separation had previously allowed us to isolate the excitatory synaptic component by iontophoretic application of the antagonist NBQX (McColgan et al., 2019), and here the inhibitory synaptic components.

The ability to separate the sources of the EFP allowed us to extract the frequency and ITD tuning of the components independently (Kuokkanen et al., 2010, 2013, 2018). In NL, the axonal contributions from ipsi- and contralateral NMs sum linearly, with phases matching the map of ITD (Kuokkanen et al., 2013). These axonal contributions are assumed to be unaffected by local block of inhibition, as supported by our observations (Fig. 8). Over the duration of the experiment, however, we observed small, opposite phase shifts of monaural neurophonic responses, and hypothesized that electrode position could drift with respect to the delay lines causing small shifts in the neurophonic ITD tuning (Fig. 8E). At each NL neuron, the neuron’s excitatory synaptic activity follows the axonal activity, with strong ITD dependent phase locking (Funabiki et al., 2011; McColgan et al., 2019). By contrast, the inhibitory synaptic components were not ITD tuned (Fig. 8F,G), consistent with the observation that most SON activity was not ITD-tuned. The last component of the EFP in NL, attributed to the postsynaptic activity of the NL neurons, is strongly ITD tuned (Kuokkanen et al., 2018). We were unable to record this component reliably using the iontophoresis electrodes, possibly because the electrode may need to be in the immediate vicinity of a single cell (Peña et al., 1996; Funabiki et al., 2011; Kuokkanen et al., 2013, 2018).

## Acknowledgments

We thank Go Ashida for helpful discussions, Sarada Viswanathan for her help with the viral tracers and Waisudin Kamal for his assistance with cell counts.

## Ethical approval and informed consent statement

All animal care and experimental procedures were conducted following NIH guidelines and approved by the University of Maryland Institutional Animal Care and Use Committee. Efforts were made to minimize animal suffering and reduce the number of animals used.

## Data availability statement

Data and scripts for reproducing figures 6-8 are available at FigShare: https://doi.org/10.6084/m9.figshare.30501755.

